# The nuclear to cytoplasmic ratio drives cellularization in the close animal relative *Sphaeroforma arctica*

**DOI:** 10.1101/2023.01.19.524795

**Authors:** Marine Olivetta, Omaya Dudin

## Abstract

The ratio of nuclear content to cytoplasmic volume (N/C ratio) is a key regulator driving maternal-to-zygotic transition in most animal embryos. Altering this ratio often impacts zygotic genome activation and deregulates the timing and outcome of embryogenesis [1–3]. Despite being ubiquitous across animals, little is known about when the N/C ratio evolved to control multicellular development. Such capacity either originated with the emergence of animal multicellularity or was co-opted from mechanisms present in unicellular organisms [4]. An effective strategy to tackle this question is to investigate close relatives of animals exhibiting life cycles with transient multicellular stages [5]. Among these are ichthyosporeans, a lineage of protists undergoing coenocytic development followed by cellularization and cell release [6–8]. During cellularization, a transient multicellular stage resembling animal epithelia is generated offering a unique opportunity to examine whether the N/C ratio regulates multicellular development. Here, we use time-lapse microscopy to characterize how the N/C ratio affects the life cycle of the best-studied ichthyosporean model, *Sphaeroforma arctica*. We uncover that the last stages of cellularization coincide with a significant increase in the N/C ratio. Increasing the N/C ratio by reducing the coenocytic volume accelerates cellularization while decreasing the N/C ratio by lowering the nuclear content halts it. Moreover, centrifugation and pharmacological inhibitor experiments suggest that the N/C ratio is locally sensed at the cortex and relies on phosphatase activity. Altogether, our results show that the N/C ratio drives cellularization in *S. arctica*, suggesting that its capacity to control multicellular development predates animal emergence.

## Results & Discussion

### Cellularization in *S. arctica* coincides with an increase in the N/C ratio

Animals are closely related to three protist lineages, choanoflagellates, filastereans and ichthyosporeans (Figure 1A) [6, 9]. Among these, ichthyosporeans exhibit a wide variety of cell behaviours, morphologies and complex life cycles [8]. Most characterized ichthyosporeans to date grow as multinucleated coenocytes in which synchronous rounds of nuclear duplication occur without cytokinesis, followed by cellularization and release [7, 8, 10, 11]. The life cycle of the model ichthyosporean *Sphaeroforma arctica* consists of three developmental stages: (i) coenocytic growth defined by timer-dependent nuclear duplication [11], (ii) arrest in coenocytic volume growth followed by formation of an actomyosin network driving plasma membrane (PM) invaginations, leading to cellularization, and (iii) release of cell-walled newborn cells (Figure 1B) [7]. During cellularization, PM invaginations lead to the formation of a transient multicellular epithelial-like cell layer [7]. This layer then detaches abruptly prior to cell release in a process termed “Flip”, splitting cellularization into “Pre-flip” and “Post-flip” stages (Figure 1B). The mechanisms allowing for such precise developmental progression in *S. arctica* are not known, but one possible pathway may involve the N/C volume ratio. Indeed, previous observations have indicated that a higher number of nuclei per volume correlated with earlier cellularization, such as for wild-type *S. arctica* growing in limited nutrient conditions [11], for *S. arctica* mutants with faster sedimentation rates [12], and *Sphaeroforma* sister species exhibiting faster nuclear divisions [12]. These observations raise the possibility that, as in the early embryo of animals, the nuclear content per cytoplasmic volume may also regulate the development of *S. arctica*. To test this hypothesis, we first measured the change in coenocytic volume and nuclear content throughout the life cycle of *S. arctica* using long-term live imaging and nuclear staining. As anticipated, we observe that coenocytes exhibit a constant growth in volume for 35.7 ± 4.2 hrs, followed by an arrest parallel to cellularization onset which persists for 6.7 ± 2 hrs until Flip (Figure 1C and 1D). After 2.7 ± 1.1 hrs, we observe a sharp decrease in volume following the release of newborn cells (Figure 1C and 1D). In parallel, using Hoechst staining, we observe that aside from a brief latency phase during which nuclear doubling time is 13.8 ± 1 hrs, the nuclear content (number of nuclei) per coenocyte increases constantly over time, with a nuclear division occurring every 8.16 ± 2 hrs until new born cells are released (Figures 1E and 1F). Notably, we observe that the number of nuclei keeps increasing during cellularization (36-42 hrs) despite the arrest in coenocytic volume growth (Figure 1C and 1F). Thus, when dividing the overall nuclear volume measured from Hoechst-stained coenocytes by the average coenocytic volume obtained from live images, we observe a doubling in the N/C ratio (~4 to ~8%) prior to Flip (Figure 1G). As this observation was inferred from averaged results of two distinct sets of experiments, we set to investigate this process at the level of single coenocytes. Using a nuclear-excluded membrane dye (FM4-64) in combination with light-sheet microscopy, we were able to track a few nuclear divisions during the last stages of coenocytic development (Video S1). We observe that the last nuclear division precedes membrane invaginations by 1.5 ± 0.9 hrs (Figure 1H and Video S1) which is significantly less than the total duration of the Pre-flip phase (Figure 1D). This result shows that the last nuclear division precedes membrane invagination and occurs during the Pre-flip phase when the coenocytic volume is constant, thereby increasing the N/C ratio (Figure 1G). To further confirm this result, we used phalloidin to stain F-actin in fixed coenocytes to differentiate between coenocytes at cellularization onset, in which actin localises as small nodes [7], and coenocytes starting PM invaginations, which exhibit actin bundles at the cortex (Figure 1I) [7]. These measurements at the level of single coenocytes confirm that the N/C ratio exhibits a significant increase at the time of PM invaginations (Figure 1J), and therefore demonstrates that the late phase of cellularization preceding Flip, coincides with an increase in the N/C ratio.

**Figure 1.**
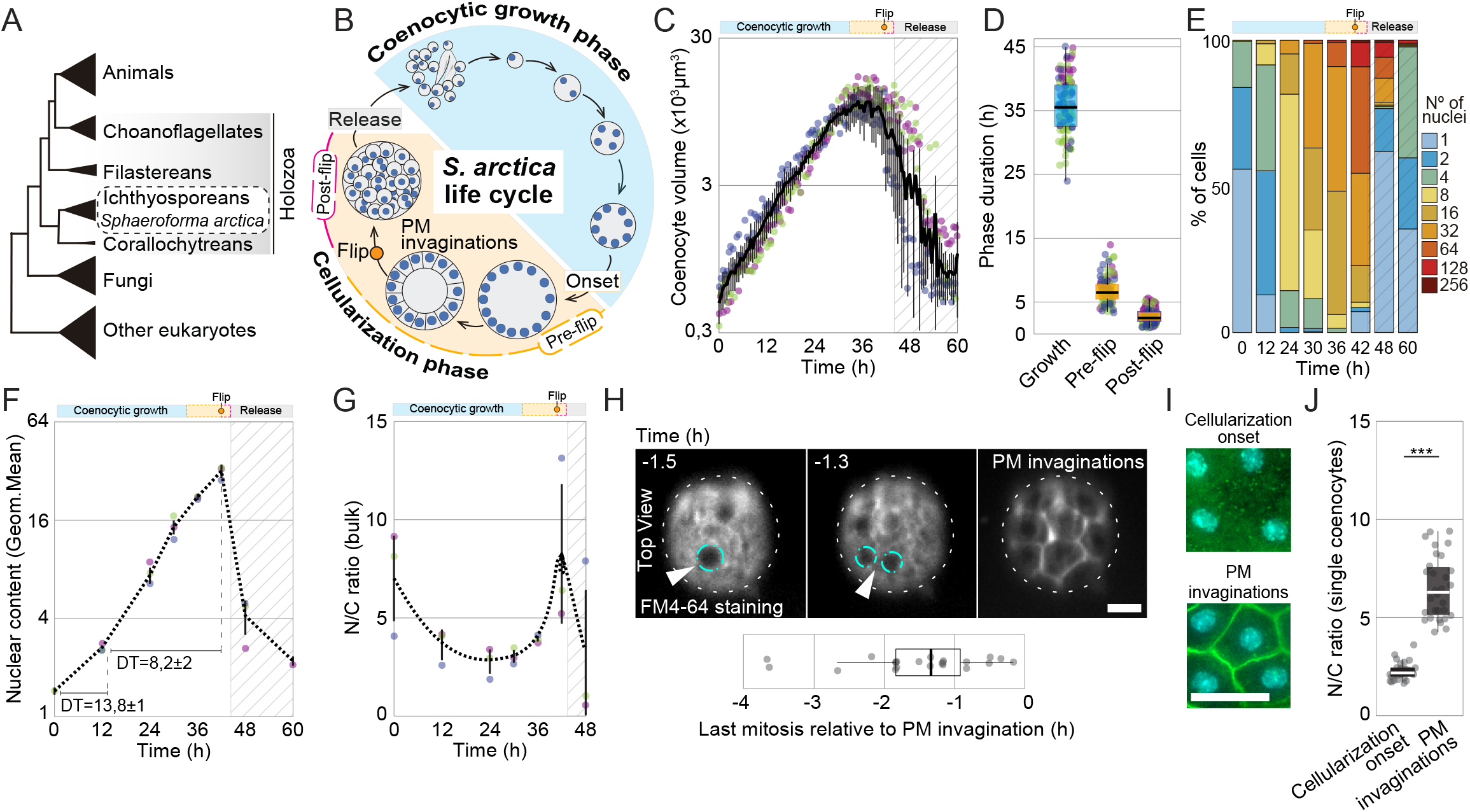
The N/C ratio increases during the latest phase of cellularization in *S. arctica*. (A) Cladogram representing the position of ichthyosporeans including *Sphaeroforma arctica* within the eukaryotic tree. (B) Schematic representation of the life cycle of *S. arctica* with coenocytic growth, Pre-flip and Post-flip as the three main developmental stages. (C) Average coenocytic volume per time point at 17°C (n >100/time point). Three independent replicates out of 6 are represented here for visibility. The black line represents the average of all 6 replicates. The grey shading indicates the point at which newborn cells are released. (D) The average duration of growth, Pre-flip and Post-flip stages at 17°C represented as box plots (n =119). (E) Distribution of nuclear content of *S. arctica* coenocytes during the life cycle at 17°C measured by microscopy of Hoechst-stained coenocytes (n >285/time point). (F) Quantification of the mean nuclear content per time point at 17°C (expressed as log2 of geometric mean) with three independent replicates visible (n >1063/time point). Doubling time (DT) measured for both latency and exponential phases are presented. (G) Average N/C ratio per time point estimated from bulk cultures. This ratio was measured by dividing the average nuclear volume obtained from Hoechst-stained coenocytes by the average coenocytic volume obtained by live microscopy. (H) Dynamics of nuclear duplications relative to PM invaginations during cellularization. Live coenocytes pre-grown for 36 hrs, were stained with FM4-64 and imaged using light-sheet microscopy. Note that the membrane dye is excluded from the nuclei (cyan dashed line) allowing to track a few nuclear divisions (arrow). Measurements of the last observed mitosis relative to membrane invaginations (T = 0) are represented in the box plot (n =22). Bar=10μm. (I) Actin cytoskeleton localisation during early and late cellularization. Synchronized coenocytes of *S. arctica* were fixed at T=42 hrs and stained with phalloidin (green) and Hoechst (cyan). Early cellularizing coenocytes exhibit small actin nodes at the cortex, whereas late cellularizing coenocytes exhibit actin bundles concomitant with PM invaginations [7]. Bar=10μm. (J) Average N/C ratio during early and late cellularization. This ratio was measured in single coenocytes by dividing the average nuclear volume by the coenocytic volume obtained (n =30/stage). The phalloidin staining allows the identification of early and late cellularizing coenocytes.

### Manipulating the N/C ratio impacts the timing and developmental outcome in *S. arctica*

To assess whether the N/C ratio controls developmental progression, we sought to alter it by either diminishing the coenocytic volume or the number of nuclei. Previous work in *S. arctica* has shown that nutrient limitation leads to a decrease in coenocytic volume without affecting nuclear divisions [11]. Thus, nutrient limitation increases the N/C ratio, allowing us to assess the consequence on cellularization timing (Figure 2A). Therefore, we monitored the N/C ratio, the timing of Flip and developmental outcome of coenocytes grown in normal media (MB), as well as various dilutions thereof (MB1/4, MB1/8, MB1/16). In agreement with our previous observations [11], we find that coenocytes growing in low nutrients undergo cellularization without altering the pace of nuclear duplication cycles (Figure 2B, 2C, S1A and S1B). Moreover, the lower the media concentration, the higher the N/C ratio is and the earlier Flip and cellularization occur (Figure 2D, S1C-S1F and Video S2). However, despite the acceleration in the timing of cellularization, the N/C ratio at the time of Flip is comparable across the different nutrient concentrations, indicating that a specific N/C ratio threshold must be reached to complete cellularization (Figure 2E). As earlier cellularization may merely reflect an indirect effect of nutrient depletion, we set out to alter the N/C ratio differently by blocking nuclear division cycles (Figure 2A). The microtubule depolymerizing agent Carbendazim (MBC) affects nuclear positioning in *S. arctica* without blocking PM invaginations or cell release when added to early cellularizing coenocytes (Figure S1G) [7]. Thus, we hypothesized that MBC may be used to block nuclear duplication at precise phases of the life cycle to decrease the N/C ratio without affecting PM invaginations *per se*. Therefore, we tracked nuclear content, coenocytic volume and development of *S. arctica* cultures treated with MBC at different time points (T0, T15, T24, T33 hrs). We find that MBC treatment prevents both nuclear divisions and subsequent cellularization from proceeding, despite having little or no effect on coenocytic volume dynamics (Figure 2F-2I, S1H and Video S2). Overall, our observations suggest that decreasing the N/C ratio by blocking nuclear division prevents cellularization. Together, these results show that in *S. arctica*, increasing the N/C ratio using starvation accelerates development whereas decreasing the N/C ratio below a certain threshold using MBC prevents developmental progression.

**Figure 2.**
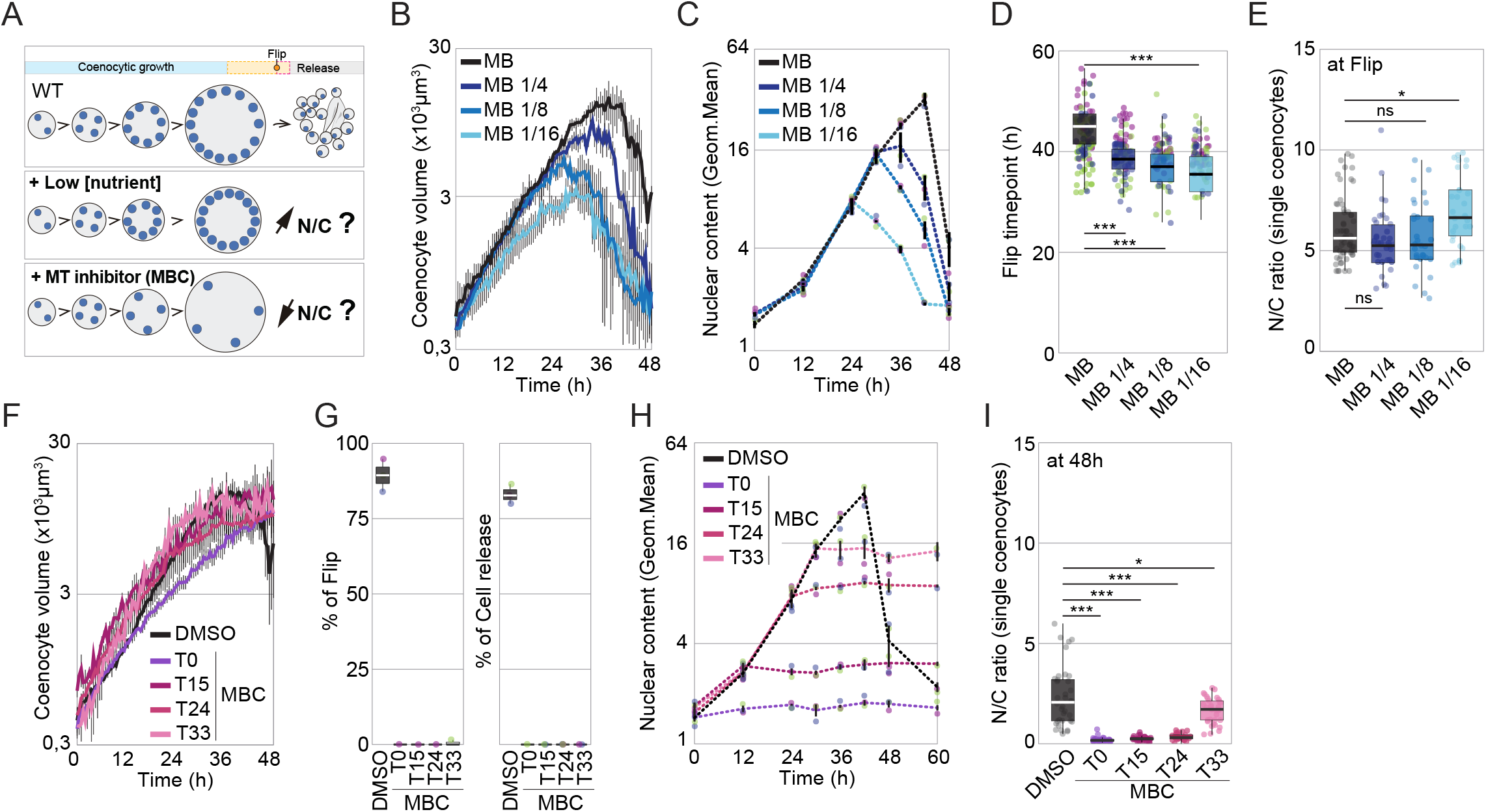
Manipulating the N/C ratio impacts the timing and outcome of *S. arctica* development. (A) Schematic representation of the two strategies used to manipulate the N/C ratio. Nutrient starvation should decrease the coenocytic volume without affecting nuclear duplication leading to an increase in the N/C ratio. MBC treatment should block nuclear duplications without affecting coenocytic volume growth leading to a decrease in the N/C ratio. (B) Average coenocytic volume per time point at 17°C for coenocytes growing in different media concentrations (n >100/time point). The different lines represent the average of at least 5 independent replicates. (C) Quantification of the mean nuclear content per time point at 17°C for coenocytes growing in different media concentrations (expressed as log2 of geometric mean) (n >209/time point). (D) Average time at which coenocytes growing in different media concentrations undergo Flip (n >200/medium). Note that coenocytes growing in lower media concentration undergo Flip earlier than the control (MB). Three independent replicates out of 6 are represented here for visibility. The box plot represents the average of all 6 replicates (*t* test, ***, P < 0.0001). (E) Average N/C ratio around Flip time point of coenocytes growing in different media concentrations (n >30/medium). This ratio was measured at T=42 hrs for coenocytes growing in MB and MB1/4 and at T=36 hrs for coenocytes growing in MB1/8 and MB1/16. Measurements were obtained from single coenocytes by dividing the average nuclear volume by the coenocytic volume. Note that the N/C ratio is similar across media concentration despite Flip occurring much earlier in MB1/8 and MB1/16. (*t* test, *, P < 0.05). (F) Average coenocytic volume per time point at 17°C for coenocytes treated with MBC at different time points of their life cycle (n >100/time point). Note that despite their incapacity to complete cellularization and release newborn coenocytes, the growth dynamics of MBC-treated coenocytes are only slightly affected when compared to the control (DMSO). The different lines represent the average of 3 independent replicates. (G) Percent of Flip and cell release of coenocytes grown in MB and treated with MBC at different time points of their life cycle (n >200/treatment). Note that MBC-treated coenocytes do not cellularize in comparison to the control (DMSO). (H) Quantification of the mean nuclear content per time point of MBC-treated coenocytes growing in MB (expressed as log2 of geometric mean) (n >200/time point). Note that the nuclear duplication cycle is consistently blocked at the time of MBC addition. (I) Average N/C ratio at T=48 hrs of MBC-treated coenocytes growing in MB (n >30/treatment). This ratio was measured in single coenocytes by dividing the average nuclear volume by the coenocytic volume. Note that the N/C ratio of MBC-treated coenocytes is significantly below 5% even at a later time point (T=48 hrs) (*t* test, *, P < 0.05, **, P < 0.005, ***, P < 0.0001).

To further investigate the role of the N/C ratio in *S. arctica*, we aimed to rescue cellularization in MBC-treated coenocytes exhibiting a low N/C ratio. We treated a culture grown for 24 hrs in normal media with MBC for 9 hrs, thus skipping a nuclear division cycle, before washing the drug away and tracking developmental outcome (Figure 3A). We observe that in comparison to coenocytes continuously treated with MBC, such blocked and released coenocytes recover about 48% of their ability to cellularize (Figures 3B and 3C). Importantly, we also find that MBC-blocked and released coenocytes exhibit an increase in the nuclear content when compared to coenocytes maintained with MBC, demonstrating that an additional nuclear duplication cycle occurred resulting in a re-increase in the N/C ratio (Figure 3D, 3E and S2A). As the additional nuclear division cycle occurs only after washing the drug away, we observe a significant delay in the increase of the N/C ratio and subsequent timing of Flip when compared to DMSO-treated coenocytes (Figures 3E-3F). Notably, we observe that DMSO-treated coenocytes undergo Flip earlier (38.8 ± 1.8 hrs) when compared to previous experiments (42.5 ± 6.3 hrs) (Figures 2D and 3F). This disparity is seemingly dependent on the experimental set-up requiring repeated centrifugation to wash out MBC (see results below). Nonetheless, our results show that delaying N/C ratio dynamics defers the timing of cellularization. These results are reminiscent of experiments demonstrating that haploid early animal embryos undergo an additional nuclear division before the mid-blastula transition (MBT) [13–16]. In a second experiment, we set out to reduce the coenocytic volume of MBC-treated coenocytes incapable of completing cellularization to ask whether a re-increase of the N/C ratio would revert this fate (Figure 3A). To do so, we combined both previous approaches by treating coenocytes with MBC at different time points of the life cycle in combination with different media concentrations (Figure 3A and 3G-3I). Our results show that coenocytes treated with MBC early on (T=0 to 15 hrs) are unable to complete development, regardless of nutrient limitation (Figures 3G and S2B-S2D). However, when MBC is added later (T=24 to 33 hrs), we observe a gradual and nutrient-dependent developmental recovery (Figures 3G, S2B-S2D and Video S2). Thus, the development of MBC-treated coenocytes can be rescued when grown in lower media concentration, which reduces overall volume and re-increases the N/C ratio (Figures 3H-3I). Our results parallel previous studies in animals, where the removal of cytoplasmic content resulted in premature MBT [16–18]. Together, our findings show that similar to early animal embryos, the timing and progression of cellularization in *S. arctica* are tightly linked and regulated by the N/C ratio.

**Figure 3.**
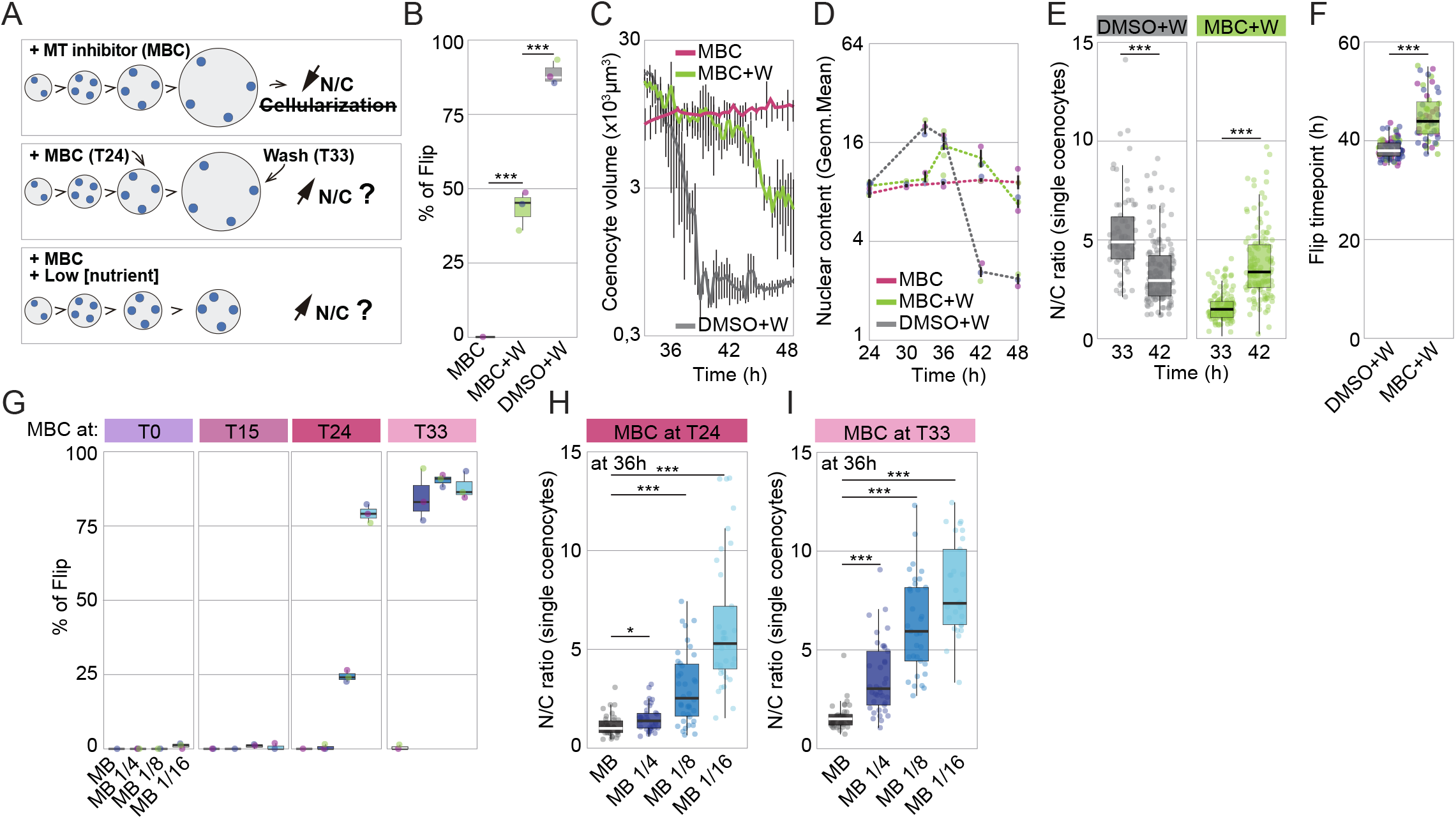
Cellularization recovery in MBC-treated coenocytes. (A) Schematic representation of the two strategies used to re-increase the N/C ratio in MBC-treated coenocytes exhibiting low N/C ratio. Adding MBC at T=24 hrs followed by a wash at T=33 hrs, also known as a block and release, should allow for renewed nuclear duplication cycles and thus a re-increase of the N/C ratio. Combining MBC treatment with nutrient starvation should decrease the coenocytic volume which may lead to a re-increase in the N/C ratio. (B) Percent of coenocytes undergoing Flip when subject to a block and release with MBC (n >100/condition). Note that following washing, MBC-treated coenocytes partly recover their capacity to complete cellularization (MBC+W) within the duration of the experiment (48 hrs) in comparison to MBC-treated coenocytes without washing (MBC) (*t* test, ***, P < 0.0001). (C) Average coenocytic volume per time point of coenocytes subject to a block and release with MBC (n >34/time point). Live measurements start at T=34 hrs to T=48 hrs. Note that cellularization of block and released coenocytes is rescued but shows a significant delay when compared to DMSO-treated coenocytes. (D) Quantification of the mean nuclear content per time point of coenocytes subject to a block and release (expressed as log2 of geometric mean). Note that following washing, MBC-treated coenocytes (MBC+W) undergo an additional nuclear duplication cycle when compared with the control (MBC) (n >158/time point). (E) Average N/C ratio at T=33 and T=42 hrs of coenocytes subject to a block and release with MBC (n >67/condition). This ratio was measured in single coenocytes by dividing the average nuclear volume by the coenocytic volume. Note that the N/C ratio of MBC-treated coenocytes re-increases following the MBC release at T=33 hrs (*t* test, ***, P < 0.0001). (F) Average time at which blocked and released coenocytes undergo Flip (n >58/condition). Note that MBC-treated and washed coenocytes (MBC+W) undergo Flip later than the control (DMSO+W) (*t* test, ***, P < 0.0001). (G) Percent of coenocytes undergoing Flip in coenocytes growing in different media concentrations and treated with MBC at different time points of their life cycle (n >100/condition). Note that MBC-treated coenocytes recover their capacity to complete cellularization when grown in lower media concentration. (H-I) Average N/C ratio at T=36 hrs of coenocytes growing in different media concentrations and treated with MBC at T=24 and T=33 hrs (n >29/condition). This ratio was measured in single coenocytes by dividing the average nuclear volume by the coenocytic volume. Note that the N/C ratio of MBC-treated coenocytes growing in lower media concentration increases when compared to the control (MB) (*t* test, *, P < 0.05, ***, P < 0.0001).

### The N/C ratio is locally sensed at the cortex during the cellularization of *S. arctica*

The N/C ratio controls cell cycle progression and zygotic gene expression during early animal development, but the specific mechanisms sensing this ratio are not yet fully understood [3, 19–27]. It has been proposed that the underlying mechanisms involve the titration of key transcriptional inhibitors and components of the DNA replication machinery, such as histones and dNTPs, by the exponential increase in nuclear content [28–36]. *In vitro* experiments using *Xenopus* egg extract identified histones H3 and H4 as key inhibitory factors titrated by the increasing concentration of DNA [31]. Similar results were also found in the fruit fly *D. melanogaster* [36], further supporting the critical role of histones in sensing the N/C ratio, be it in early cleaving embryos like *Xenopus* or syncytial ones like *Drosophila*. Moreover, recent work suggested that the N/C ratio is not sensed globally at the level of the embryo in *D. melanogaster*, but rather locally at the level of a community of nuclei at the syncytial embryo cortex [37, 38]. This process is partly regulated by the activity of mitotic phosphatases including Protein phosphatase 1 (PP1) [27]. Localized activation of PP1, which spreads from the nuclei to the embryo cortex, activates cortical actomyosin in a spatially restricted manner. This leads to a localized increase in myosin II gradients, driving both cortical and cytoplasmic flows and ultimately ensuring efficient cellularization [24, 27, 37]. Given that several above-mentioned regulators have been identified in the genome of *S. arctica* [7, 39–41], we wondered whether similar pathways may be involved in sensing the N/C ratio. Due to the current limitations in genetic tools for ichthyosporeans, we implemented centrifugal displacement to assess, in an orthogonal manner, whether the N/C ratio is sensed globally or locally at the cortex [42–45]. We hypothesized that centrifugation should displace nuclei, leading to a local increase in cortical nuclear density (Figure 4A). If the N/C ratio is sensed globally at the level of the entire coenocyte, then centrifugation should not alter developmental timing. In contrast, if the N/C ratio is sensed locally, then centrifuged coenocytes with tighter nuclear distribution at the cortex may undergo Flip and complete cellularization earlier. To assess nuclear density, we measured the average distance from each nucleus to its nearest neighbour. We find that the distance between nuclei decreases during membrane invaginations when compared to cellularization onset, indicating a rise in cortical nuclear density (Figure 4B). Additionally, this distance remains comparable across various nutrient concentrations at the time of Flip (Figure S3A) and notably increases when treated with MBC (Figure S3B), making it a reliable indicator for nuclear density at the cortex. Using this indicator in conjunction with centrifugal displacement experiments, we can evaluate whether the N/C ratio is sensed globally or locally in *S. arctica*. For this, we used coenocytes entering the cellularization phase (T=36 hrs) to ensure they contain factors required for cellularization onset, subjected them to centrifugation, and monitored developmental outcome (Figure 4A). We observe that 60% of coenocytes exhibit an asymmetrical shift in nuclear localisation and an increase in cortical nuclear density, as evident from the shortened distance between nuclei, without changing the overall N/C ratio (Figures 4C-4F and S3C). By combining centrifugation with live imaging of the PM, we note that most centrifuged coenocytes undergo Flip and release, but ~10% undergo lysis and ~40% exhibit irregular PM invaginations (Figures 4G-4I, S3D and S3E). Indeed, PM invaginations occur asymmetrically from one side of the coenocytes, leading to the release of abnormally-sized cells at the opposite end (Figure 4H, Video S3). Such irregular cellularization was associated with earlier Flip (Figure 4J), indicating that centrifuged coenocytes undergo PM invaginations prematurely in response to the local increase in nuclear density. Together, these results suggests that the N/C ratio is sensed locally at the cortex. To explore the underlying mechanisms of N/C sensing, we tested the impact of small molecule inhibitors on centrifuged coenocytes exhibiting increased nuclear density (at T=36 hrs). We initially examined if microtubules are necessary for sensing the N/C ratio using MBC. As previously determined, we anticipated that MBC treatment would not prevent PM invaginations when the appropriate N/C ratio is reached (Figures 3G-2I) [7]. As predicted, we observe no effects with MBC treatment, indicating that sensing the N/C ratio at the cortex is independent of microtubules (Figures 4K, 4L, and S3F-S3H). In order to determine if the N/C ratio sensing is dependent on a signal originating from within the nuclei, we inhibited both the nuclear export using Leptomycin B (LTB) and the nucleocytoplasmic transport using Wheat Germ agglutinin (WGA). Our results show no significant impact on developmental timing or outcome, suggesting that N/C ratio sensing at that stage relies on a cytoplasmic signal (Figures 4K, 4L, and S3F-S3H). Lastly, to explore the role of protein phosphatases, we applied a broad phosphatase inhibitor cocktail (PanPi) and a specific serine/threonine phosphatase inhibitor Calyculin A (CalyA) that targets both PP2A and PP1. Our results show that PanPI- and CalyA-treated coenocytes display increased lysis and a significant decrease in cellularization efficiency (complete block with PanPI), indicating that cellularization relies on efficient phosphatases activity, possibly PP1 (Figures 4K, 4L, and S3F-S3H). Although pleiotropic effects for the tested inhibitors cannot be ruled out, our data suggests that the N/C ratio is locally sensed through a cytoplasmic signal requiring phosphatase activity.

**Figure 4.**
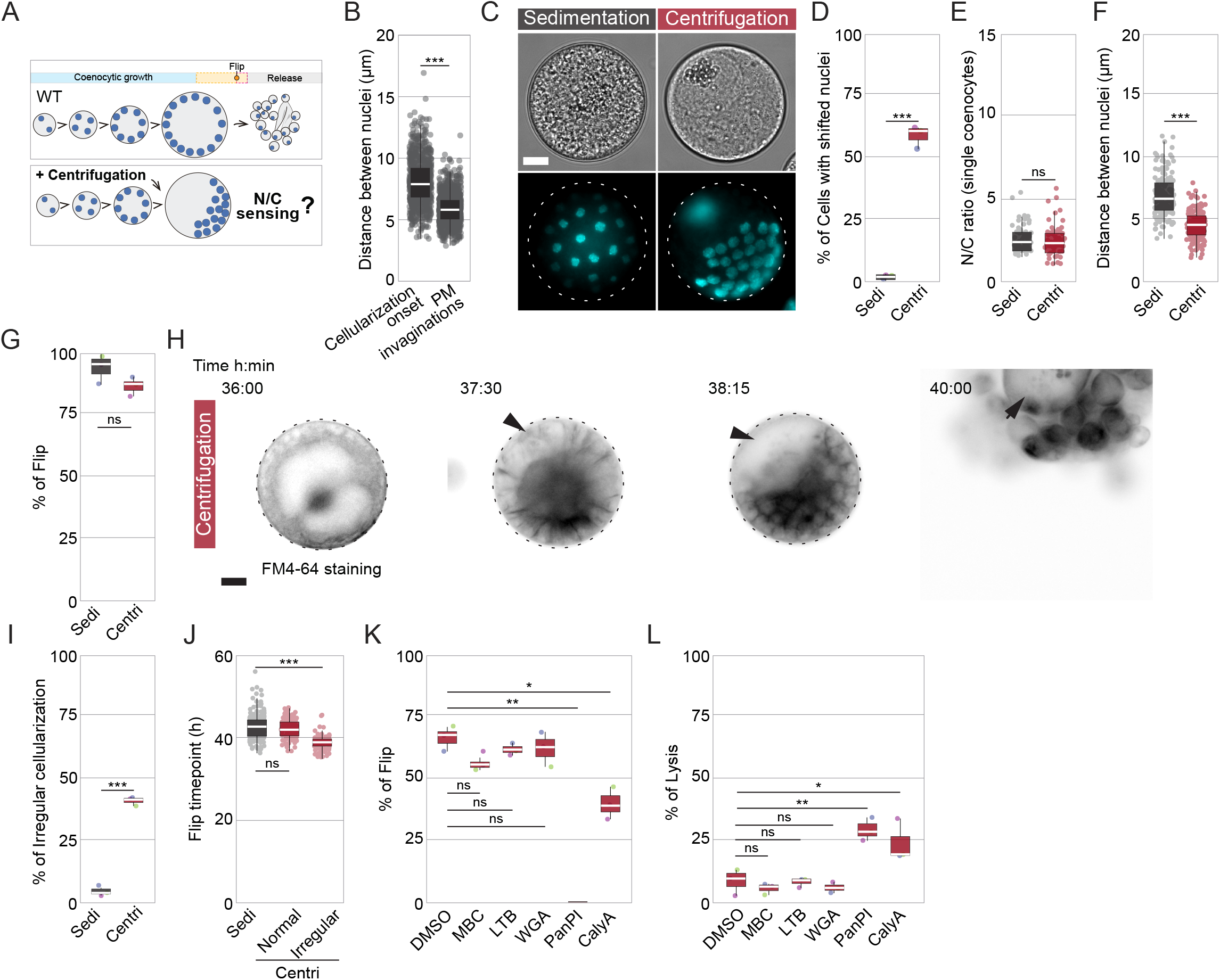
Local increase of cortical nuclear density triggers premature cellularization. (A) This scheme illustrates our strategy to uncover the mechanisms of N/C ratio sensing. By subjecting the coenocytes to centrifugation, the localisation of the nuclei will shift, leading to a localised increase in nuclear density at the cortex without altering the overall N/C ratio. This can be used to determine whether the N/C ratio is sensed locally at the cortex or globally throughout the coenocyte. (B) Average distance between nuclei at the cortex during early and late cellularization (n >800/stage). Note that nuclei are closer to each other parallel to PM invaginations reflecting an increased nuclear density at the cortex before Flip (*t* test, ***, P < 0.0001). (C) Coenocytes pre-grown for 36 hrs were either left to sediment or centrifuged at 12000 rpm for 5 min before fixation and Hoechst staining. Note that centrifuged coenocytes exhibit a shift in nuclear localisation on one side, with a few lipidic and auto-fluorescent granules localised on the opposite side. Bar=10μm. (D) Percent of coenocytes exhibiting shifted nuclear localisation after sedimentation (Sedi) or centrifugation (Centri) (n >95/condition) (*t* test, ***, P < 0.0001). (E) Average N/C ratio in sedimented and centrifuged coenocytes at T=36 hrs (n >50/condition). This ratio was measured in single coenocytes by dividing the average nuclear volume by the coenocytic volume obtained. Note that the N/C ratio is not altered following centrifugation (Centri). (F) Average distance between nuclei at the cortex in sedimented and centrifuged coenocytes at T=36 hrs (n >152/condition). Note that nuclei are closer to each other in centrifuged coenocytes (Centri) reflecting an increased nuclear density at the cortex in comparison to sedimented coenocytes (Sedi) (*t* test, ***, P < 0.0001). (G) Percent of coenocytes undergoing Flip after sedimentation or centrifugation (n >199/condition). (H) Time-lapse images of centrifuged and FM4-64-stained coenocytes. Note that membrane invaginations occur asymmetrically and cellularization is completed irregularly with an abnormally-sized newborn cell released (arrow). Bar=10μm. (I) Percent of coenocytes undergoing irregular cellularization after sedimentation or centrifugation (n >199/condition). (J) The average time at which sedimented and centrifuged coenocytes undergo Flip (n >91/condition). Note that centrifuged coenocytes exhibiting irregular cellularization undergo Flip significantly earlier than the control (Sedi) (*t* test, ***, P < 0.0001). (K) Percent of centrifuged coenocytes undergoing Flip after treatment at T=36hrs with small molecule inhibitors targeting key cellular pathways (n >97/condition). Carbendazim (MBC) inhibits microtubule polymerization and alters cortical nuclear localisation, leptomycin B (LTB) is a nuclear export inhibitor, Wheat Germ Agglutinin (WGA) inhibits nucleocytoplasmic transport, Phosphatase inhibitor cocktail I (PanPi) is a broad phosphatase inhibitor and Calyculin A (CalyA) is a serine/threonine phosphatase inhibitor such as PP1 and PP2A (*t* test, *, P < 0.05, **, P < 0.005). (L) Percent of centrifuged coenocytes undergoing lysis after treatment with small molecule inhibitors targeting key cellular pathways (n >97/condition) (*t* test, *, P < 0.05, **, P < 0.005).

In summary, we show that the development of the ichthyosporean *S. arctica*, a close animal relative, is regulated by the N/C ratio. Manipulating the N/C ratio by altering either coenocytic size or nuclear content impacts developmental timing and outcome. Our results using centrifugation suggest that the N/C ratio is sensed locally at the cortex and involves phosphatase activity. While the underlying molecular mechanisms are yet to be uncovered, our findings suggest that the role of the N/C ratio in driving cellularization in *S. arctica* and, consequently, its transition to multicellularity, is reminiscent of N/C ratio-dependent regulation of early animal embryogenesis. This work positions *S. arctica* as a promising model organism for understanding the evolutionary origins and pre-metazoan mechanisms of N/C ratio-dependent regulation of multicellular development.

## Acknowledgements

We thank Pierre Gönczy, Gautam Dey, Hiral Shah and Margarida Araujo for critical reading of the manuscript; Andy Oates, Petr Strnad and Andrea Boni for the access to the LS1 Live light sheet microscope; and all the members of the Pierre Gönczy lab for helpful discussions and technical advice. This work was funded by an Ambizione fellowship from the Swiss National Science Foundation (PZ00P3_185859). The funders had no role in study design, data collection and analysis, decision to publish, or preparation of the manuscript.

## Author contributions

Conceptualization, O.D.; Methodology, O.D.; Investigation, M.O., and O.D.; Writing – Original Draft, O.D.; Writing – Review & Editing, M.O., and O.D.; Funding Acquisition, O.D.; Supervision, O.D.

## STAR METHODS

### Experimental model and subject details

The experimental model used is the ichthyosporean *Sphaeroforma arctica* originally described in [46]. A frozen *S. arctica* culture cryopreserved in 2012 at −80°C was recently diluted and maintained in Marine Broth 2216 (MB) (BD Difco 279110, or Sigma 76448, 37.4 g/L) at 17°C. For media composition experiments, MB was diluted to the desired concentration with artificial seawater (Instant Ocean, 36 g/L).

### Method details

#### Culturing conditions

*S. arctica* cultures were grown and synchronized as described previously [7, 11]. Briefly, saturated cultures in MB were diluted into a fresh medium at low density (1:200 dilution of the saturated culture) and grown in rectangular canted neck cell culture flasks with vented cap (Falcon; ref: 353108) at 17°C in the dark, resulting in a synchronously growing culture. Saturated cultures of *Sphaeroforma* sp. are obtained after 3 weeks of growth in MB or ~5 days of growth in MB 1/16 diluted in with artificial seawater. Various dilutions of MB were used throughout this study (MB1/4, MB1/8 and MB1/16), all of which were done by diluting MB with artificial seawater.

#### Microscopy

Microscopy of live and fixed coenocytes was performed using a fully motorized Nikon Ti2-E epifluorescence inverted microscope equipped with a hardware autofocus PFS4 system, a Lumencor SOLA SMII illumination system and a Hamamatsu ORCA-spark Digital CMOS camera. A CFI Plan Fluor 20X, 0.50 NA., a CFI Plan Fluor 40X Air and a CFI Plan Fluor 60X Oil, 0.5-1.25 NA. objectives were used for imaging. To maintain the temperature at 17°C we used a cooling/heating P Lab-Tek S1 insert (Pecon GmbH) connected to a Lauda Loop 100 circulating water bath. Light sheet microscopy in (Figure 1H, Video S1) was performed using the LS1 Live light sheet microscope system (Viventis®) previously described in detail in [47, 48] using a 25X 1.1 NA objective (CFI75 Apo 25XW; Nikon) and an sCMOS camera (Zyla 4.1, Andor). Light sheet imaging was conducted in a room specifically cooled at 17°C using an air-conditioning unit.

#### Cell fixation and staining

Cell fixation was performed using 4% formaldehyde and 250mM Sorbitol for 20 minutes before being washed twice with PBS. For nuclei staining, cultures were allowed to sediment for 15 minutes at RT before fixation and addition of Hoechst 33342 nuclear stain (ThermoFisher; ref: 62249) at 20μM. For actin staining, fixed coenocytes were washed once with PBS before the addition of the F-actin stain Alexa Fluor™ 488-Phalloidin (ThermoFisher; ref: A12379) at a final concentration of 0.165μM. Before imaging, fixed and stained coenocytes were concentrated before being disposed between slide and coverslip. Hoechst-stained samples were imaged to measure the nuclear content, coenocyte size and N/C ratio.

#### Live microscopy

For live-cell imaging saturated cultures were diluted 200X in various marine broth diluted media inside an 8-well ibidi chamber (Ibidi; ref:80826). To ensure oxygenation during the whole period of the experiment, the plastic cover was removed. To reduce light toxicity, we used a 715nm LongPass Color Filter (Thorlabs; ref: FGL715S). For plasma membrane live staining (Figures 1H and 4H, Videos S1 and S3), FM4-64 (ThermoFisher; ref: T13320) at a final concentration of 10μM from 100X DMSO diluted stock solution was directly added to the medium before imaging.

#### Small molecule inhibitors and centrifugal displacement

Treatment with small molecule inhibitors was performed on live coenocytes inside an 8-well chamber (Ibidi). Carbendazim (MBC) (Sigma-Aldrich; ref: 378674) was used at a final concentration of 25μg/ml from a stock of 2.5mg/ml in dimethyl sulfoxide (DMSO). Leptomycin B (LTB) (eMolecules Inc; ref: 293138671) was diluted 1000X from a stock at 2000nM in Ethanol. Wheat Germ Agglutinin (WGA) (Chemie Brunschwig AG; ref: BIO29021) was used at a final concentration of 2nM from a stock of 4μM in Water. Phosphatase inhibitor cocktail Set I (PanPi) (Lucerna-Chem AG; ref: RAY-AA-PHI-I-1) was diluted 1000X from a stock of 100X in Water. Calyculin A (CalyA) (Sigma-Aldrich; ref: 5082260001) was used at a final concentration of 50nM from a stock of 10μM in DMSO. For blocked and released coenocytes, pre-grown and drug or control-treated *S. arctica* (33 hrs) were washed three times in consumed medium (Medium in which *S. arctica* coenocytes have been growing in parallel to ensure the least perturbations) using centrifugation steps (3000 rpm for 5min) before imaging as indicated previously. For experiments involving MBC treatments in presence of various MB growth media, coenocytes were treated with MBC at 25μg/ml at the indicated growth time by exogenous addition in the well or flask. Centrifugal displacement was performed on pre-grown coenocytes (36 hrs) using 12000 rpm for 5 minutes, after which cell fixation or live-imaging with or without the addition of small molecule inhibitors was performed as previously indicated.

#### Image analysis

Image analysis was done using ImageJ software (version 1.52) [49]. For nuclear content, distribution fixed and Hoechst-stained coenocytes were imaged, and the number of nuclei per coenocyte was counted using the ObjectJ plugin in ImageJ. To compute nuclear duplication times, log2 of the geometric mean of the nuclear content was calculated as: log_2_(geommean) = ∑*_i_f_i_*\* log_2_ (*x_i_*) where fi is the fraction of coenocytes and xi the nuclear content (number of nuclei per cell) (ploidy) of each i-th nuclear content bin. Nuclear doubling times were computed as the linear regression of log2 of the geometric mean of nuclear content versus time. For measurements of the coenocytic volume in live imaged movies, images were transformed into a binary before using the particle analysis function in ImageJ with a circularity parameter set to 0.15 to 1 to measure the perimeter of the coenocyte. Values were generated as an average of all present coenocytes. For measurements in fixed coenocytes, we used the oval selection tool to draw the contour of each coenocyte and measured the perimeter. As coenocytes are spherical, we computed the coenocytic volume as *V* = 4/3*πr^3^* where *r* is the coenocyte radius. For bulk measurements of nuclear to cytoplasm ratio, we multiplied the geometrical mean of nuclear content (see above) by the average volume observed per nuclei measured from >100 independent interphase nuclei (20.77±3.03μm^3^) to generate an averaged nuclear volume, divided it by the average coenocytic volume obtained from live-imaged movies (see above) and represented in percentage. For single coenocyte measurements of the nuclear to cytoplasm ratio, we multiplied the number of nuclei observed per coenocyte by the average volume observed per nucleus (see above), divided it by the coenocytic volume measured from the same cell (see above) and represented in percentage. All experiments were performed a minimum of three independent times. For distance measurements between the closest nuclei, a line was drawn from the centre of the nuclei to their nearest neighbours. All figures were assembled with Illustrator CC 2020 (Adobe®). Several figures were generated using ggplot2 in *R* version 4.0.5 [50]. All error bars are standard deviations of at least three independent experiments except for Figures 1H, 1J, 2E, 2I, S1F, 3E, 3H, 3I, 4B, 4E, 4F, 4J, S3A, S3B, S3H in which the error bars are the standard deviation of the number of indicated samples analyzed.

---

**Figure S1.**
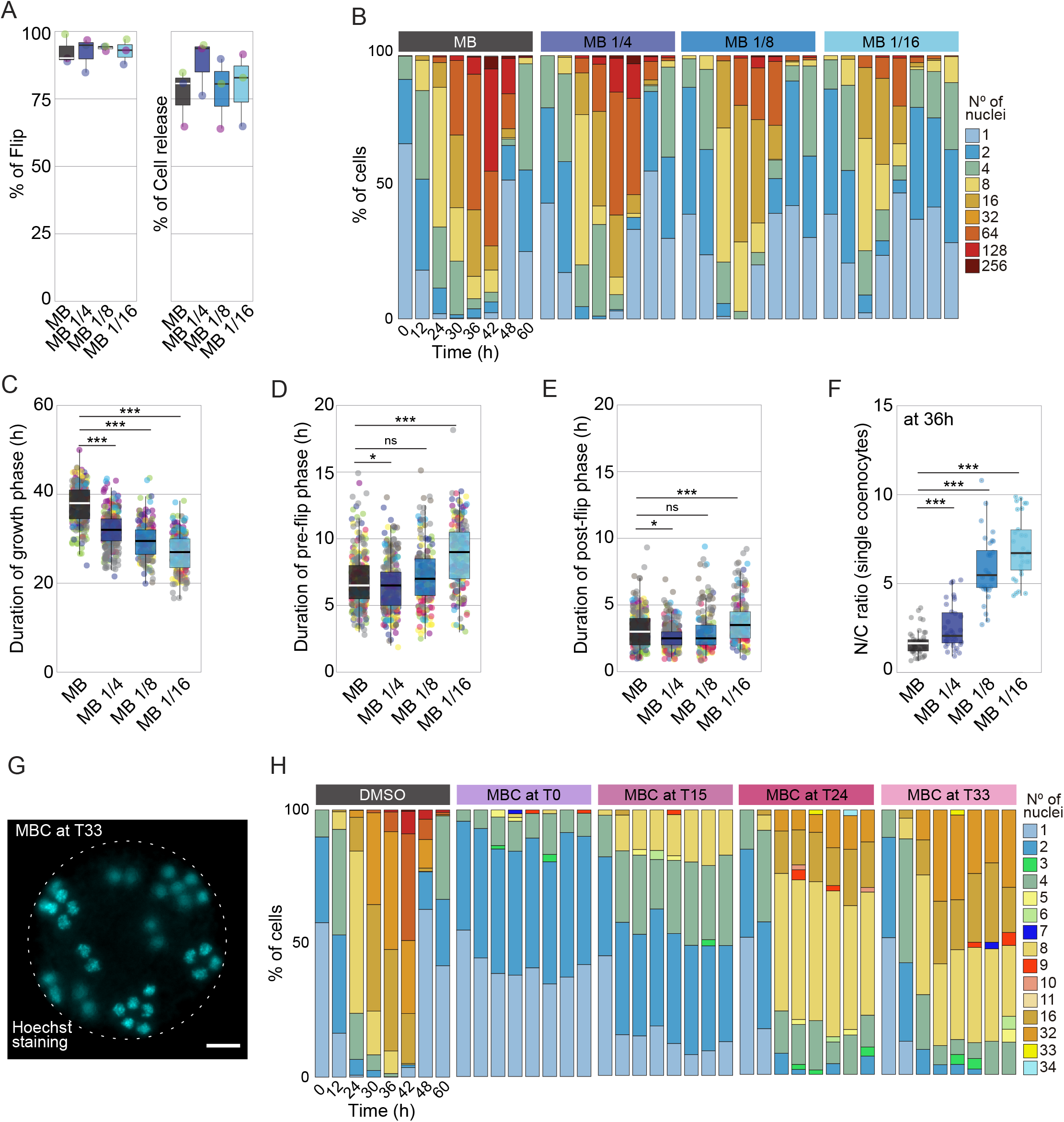
Dynamics of the N/C ratio in coenocytes grown in different media concentrations or treated with a microtubule depolymerizing agent. (A) Percent of coenocytes undergoing Flip and cell release grown in different media concentrations (n >200/medium). (B) Distribution of nuclear content of coenocytes growing in different media concentrations measured by microscopy of Hoechst-stained coenocytes (n >209/time point). Note that uninucleated newborn cells (light blue) arise earlier in lower media concentration (MB1/16) when compared to control (MB). (C-E) The average duration of growth, Pre-flip and Post-flip stages of coenocytes growing in different media concentrations (n >200/medium). Note that coenocytes growing in lower media concentrations (MB1/16) enter and undergo cellularization earlier than the control (MB). Six independent replicates are represented (*t* test, *, P < 0.05, ***, P < 0.0001). (F) Average N/C ratio at T=36 hrs of coenocytes growing in different media concentrations (n >30/medium). This ratio was measured in single coenocytes by dividing the average nuclear volume by the coenocytic volume. Note that the N/C ratio of coenocytes growing in lower media concentration (1/16MB) is significantly higher beyond 5% when compared to the control (MB) (*t* test, ***, P < 0.0001). (G) Nuclear staining using Hoechst of coenocytes treated with MBC at T=33 hrs depicting mislocalisation of nuclei. Bar=10μm. (H) Distribution of nuclear content of coenocytes growing in MB and treated with MBC at different time points of their life cycle measured by microscopy of Hoechst-stained coenocytes (n >200/time point). Note that the nuclear duplication cycle is consistently blocked at the time of MBC addition. Also, note that some coenocytes exhibit odd numbers of nuclei due to microtubule depolymerization at mitosis.

**Figure S2.**
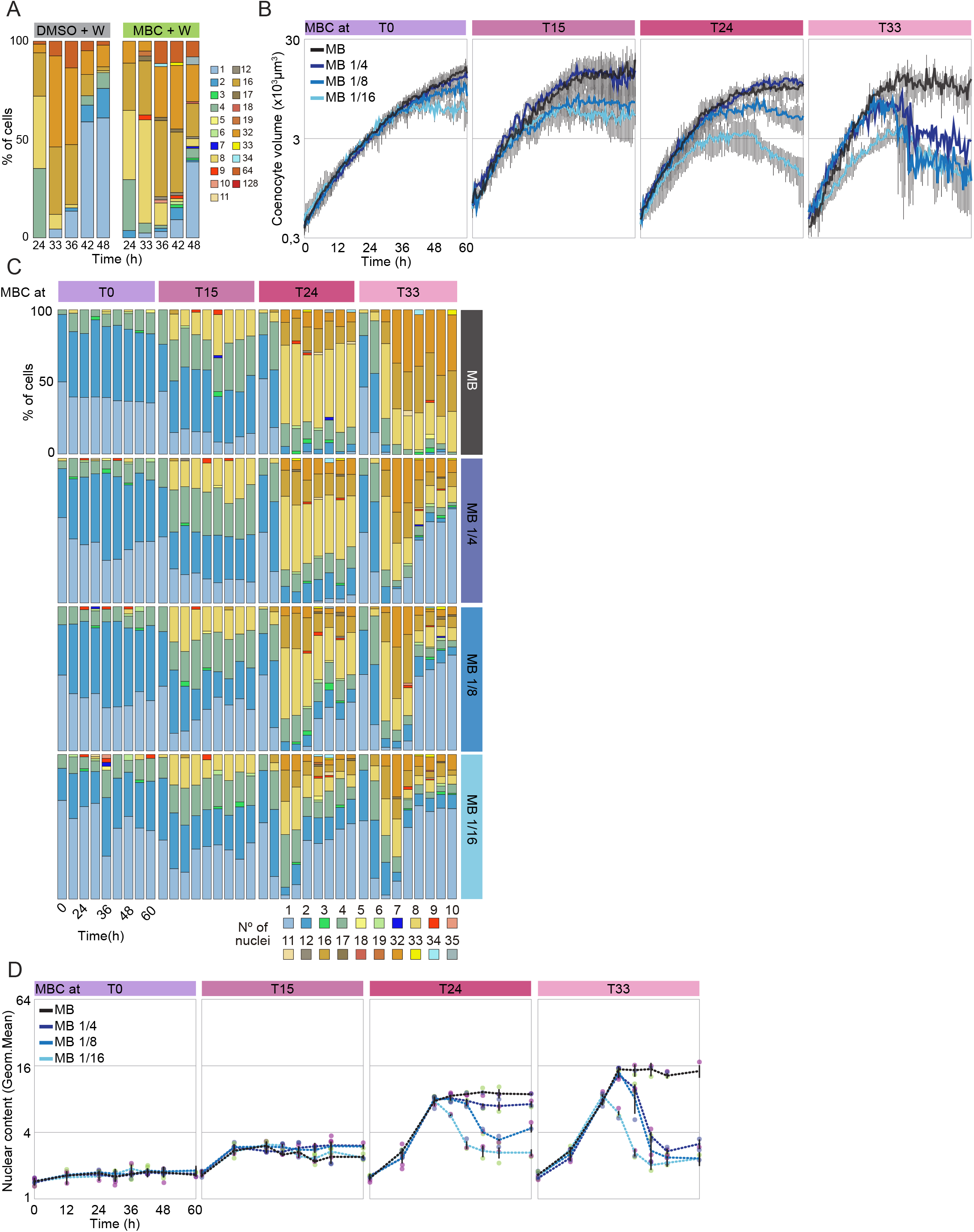
Dynamics of the N/C ratio in MBC-treated coenocytes when subject to a block and release and when grown in different media concentrations. (A) Distribution of nuclear content of coenocytes subject to a block and release with MBC (n >200/time point). Note that MBC-treated coenocytes recover their capacity to release uninucleated newborn coenocytes (light-blue) following MBC release and this recovery is delayed when compared to the control (DMSO+W). Also, note that some coenocytes exhibit odd numbers of nuclei due to microtubule depolymerization at mitosis. (B) Average coenocytic volume per time point of MBC-treated coenocytes grown in different media concentrations (n >48/time point). (C) Distribution of nuclear content of MBC-treated coenocytes grown in different media concentrations (n >100/time point). (D) Quantification of the mean nuclear content per time point of MBC-treated coenocytes grown in different media concentrations (expressed as log2 of geometric mean) (n >100/time point).

**Figure S3.**
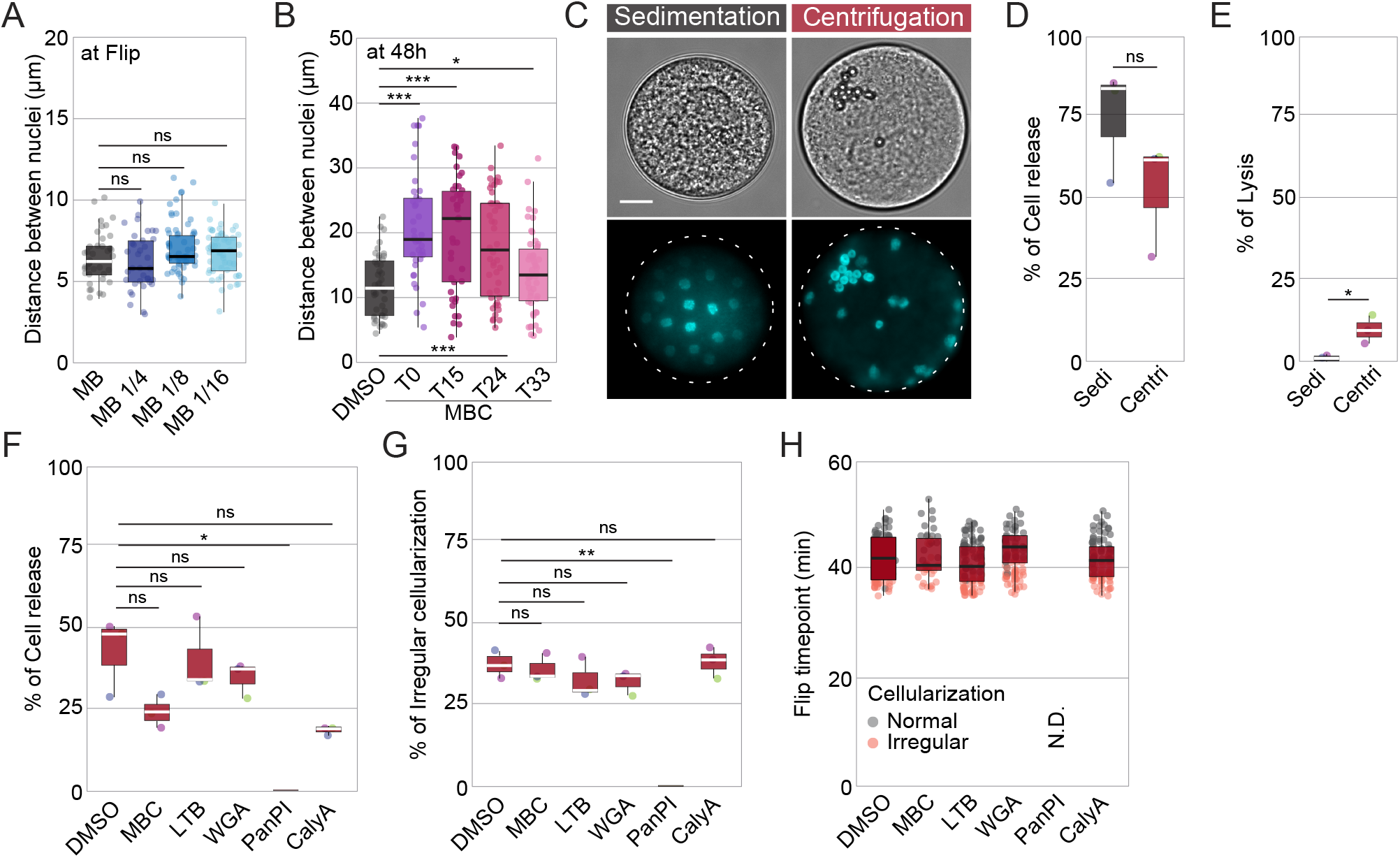
Developmental outcome of centrifuged and drug-treated coenocytes. (A) Average distance between nuclei around Flip time point of coenocytes growing in different media concentrations (n >30/medium). This distance was measured at T=42 hrs for coenocytes growing in MB and MB1/4 and at T=36 hrs for coenocytes growing in MB1/8 and MB1/16. Note that the distance between nuclei is similar across media concentration despite Flip occurring much earlier in MB1/8 and MB1/16. (B) Average distance between nuclei of MBC-treated coenocytes growing in MB shows that nuclear density is decreased compared to the control (DMSO), even at a later time point (T=48 hrs). Notably, the broader distribution of distances between nuclei in MBC-treated coenocytes is due to nuclear mispositioning and clustering (Figure S1G) (*t* test, *, P < 0.05, ***, P < 0.0001). (C) Additional example of *S. arctica* coenocytes pre-grown for 36 hrs were either left to sediment or centrifuged at 12000 rpm for 5 min before fixation and Hoechst staining. Bar=10μm. (D) Percent of coenocytes undergoing cell release after sedimentation or centrifugation (n >199/condition). (E) Percent of coenocytes undergoing lysis after sedimentation or centrifugation (n >199/condition) (*t* test, *, P < 0.05). (F) Percent of centrifuged coenocytes undergoing cell release after treatment with small molecule inhibitors targeting key cellular pathways (n >97/condition) (*t* test, *, P < 0.05). (G) Percent of centrifuged coenocytes undergoing irregular cellularization after treatment with small molecule inhibitors targeting key cellular pathways (n >97/condition) (*t* test, **, P < 0.005). (H) The average time at which centrifuged and drug-treated coenocytes undergo Flip (n >97/other conditions).

**Video S1. Time-lapse of synchronized *S. arctica* coenocytes stained with the nuclear-excluded membrane dye FM4-64 and imaged using light-sheet microscopy.**

Time interval between frames is 1min. The movie is played at 7fps. The left panel shows a top view while the right panel shows a middle section. Note that the last nuclear duplication cycle occurs approximately 1hr before PM invaginations are perceived. Bar =10μm.

**Video S2. Time-lapse of synchronized *S. arctica* coenocytes growing in different media concentrations and treated with MBC at different time points of their life cycle obtained with bright-field microscopy.** Time interval between frames is 15mins. The movies are played at 7fps. From left to right: MB, MB1/4, MB1/8, MB 1/16. From top to bottom: DMSO, MBC at T=0, MBC at T=15, MBC at T=24, MBC at T=33. Note that untreated coenocytes (in DMSO) growing in lower media concentration undergo Flip earlier than the control (MB). Moreover, early MBC treatment at T=0 prevents cellularization independent of nutrient concentrations. Lastly, MBC-treated coenocytes (T=15 to T=33) recover their capacity to complete cellularization when grown in lower media. Bar = 50μm

**Video S3. Time-lapse of centrifuged and membrane-stained *S. arctica* coenocytes undergoing irregular cellularization.** Time interval between frames is 15mins. The movies are played at 7fps. Note that membrane invaginations occur asymmetrically and cellularization is completed irregularly with an abnormally-sized newborn cell released. Bar = 10μm

